# A Rapid Gene Expression Profiler Classifies AML Tumor Responsiveness to Standard Therapies

**DOI:** 10.1101/2025.10.08.681215

**Authors:** Stephen E. Kurtz, Christopher A. Eide, Andy Kaempf, Nicola Long, Alexandria Miller, Daniel Bottomly, Shannon K. McWeeney, Curtis Lachowiez, Sudharshan Anand, Stanley W.K. Ng, Jean C.Y. Wang, John E. Dick, Jeffrey W. Tyner

**Author notes:** Correspondence: Jeffrey W. Tyner, PhD. Equal contribution (Stephen E. Kurtz, Christopher A. Eide).

## Abstract

The emergence of transcriptional signatures that define cell types and pathways has made it possible to guide cancer therapy selection through gene expression profiling. We developed a rapid qPCR-based platform to profile cell state, stemness, and BCL2 family gene expression as a companion diagnostic test for acute myeloid leukemia (AML). We validated the stability and utility of the signatures across multiple measurement platforms and using patient samples from two centers. Integrating these signatures with clinical features enables an expedient means to predict the likelihood of patient responses to two standard-of-care therapies: intensive chemotherapy and hypomethylating agent plus venetoclax (HMA+Ven). For patients treated with HMA+Ven, expression levels of the promonocyte-like signature and BCL2 add predictive value for response and overall survival in multivariable models that include genetic features. The incorporation of the rapid profiler into the prospective evaluation of newly diagnosed AML patients may enhance treatment stratification and improve outcomes.

## Introduction

Acute myeloid leukemia (AML) is characterized by considerable genetic heterogeneity resulting from a variety of mutations and chromosomal translocations that contribute to disease and influence depth and duration of responsiveness to treatments^1^. The standard-of-care for fit patients with minimal comorbidities is intensive chemotherapy induction followed by either maintenance therapy or hematopoietic stem cell transplantation. However, not all eligible patients respond to intensive chemotherapy; complete remission (CR) is achieved in 60-80% of patients <60 years and 40-60% in patients >60 years^2^. For older and less fit patients ineligible for intensive chemotherapy, standard-of-care treatment, which previously was a hypomethylating agent, such as azacitidine (Aza), now incorporates venetoclax, a specific BCL2 inhibitor^3,4^, in combination (HMA+Ven)^5^. Although response rates with HMA+Ven induction treatment in newly diagnosed (ND) elderly or frail AML patients are nearly 70%, median progression-free and overall survival, while still representing improvements over Aza alone, are approximately 10 and 15 months, respectively^5^. These outcomes highlight the importance of both defining the underlying features of patients who best respond to treatment and identifying alternative therapeutic strategies to manage emergent resistance to therapy.

AML classification based on the European LeukemiaNet (ELN) 2022 guidelines is useful in predicting overall survival (OS) outcomes for AML patients treated with intensive chemotherapy^6^. These categories are principally based upon the presence of certain cytogenetic and mutational features. However, the prognostic value for these risk categories is not retained for patients treated with HMA+Ven. A recently described molecular signature (mPRS) based on mutation status stratified outcomes for HMA+Ven treatment across three groups: low benefit (*TP53* mutated), intermediate (FLT3-ITD or N/KRAS mutated), and high benefit (all other abnormalities) with differences in median OS, which was validated in an independent cohort ^7^. These findings led to new risk classifications for patients treated with less intensive regimens^8^.

The incorporation of gene expression profiles has the potential to further improve prognostic stratification. In patients treated with intensive chemotherapy, the persistence of leukemic stem cells associates with refractoriness and relapse^9^. Gene expression profiles based on functionally-validated leukemia stem cell populations led to the definition of a 17-gene signature (LSC17) that predicts survival outcomes and responsiveness to standard induction (i.e., “7+3”) chemotherapy^10,11^. Patients with high LSC17 scores have a lower chance of achieving CR with standard induction chemotherapy and shorter OS. More recent gene expression profiling has enabled classification of AML tumors across a broader spectrum of differentiation states^12^. Among these AML cell states, progenitor-like leukemic cells have been shown to be sensitive to HMA+Ven due to high BCL2 expression levels, whereas, more differentiated AML tumors (e.g. monocyte-like) exhibit reduced HMA+Ven sensitivity^13-23^. In addition, these AML cell state signatures have been aligned with ex vivo response to a broader panel of approved and exploratory drugs and the majority of drugs exhibited sensitivity and/or resistance that correlated with AML tumor cell state^19,20,22^. These findings support incorporating gene expression profiles as a companion diagnostic to refine predicted benefit from either standard 7+3 chemotherapy or HMA+Ven, while preempting overtreatment and, consequently, reducing toxicity and costs. This companion diagnostic may also aid in predicting benefit to other cell state -governed therapeutics as they become available for use in AML. We, therefore, developed a rapid gene expression-based method, termed Myelo-ID, to simultaneously profile AML stemness, tumor differentiation states, and target gene expression.

## Methods

### Study approval

Clinical specimens were collected following informed consent from patients according to a protocol approved by the Institutional Review Board at Oregon Health & Science University (OHSU) (IRB# 4422). Biological samples from Princess Margaret Cancer Centre, Toronto, Canada (PM) were collected with informed consent according to procedures approved by the Research Ethics Board and viably frozen in the PM Leukemia Bank. All research was conducted in accordance with the Declaration of Helsinki.

### Patient populations

#### Newly Diagnosed AML Reference cohort

Samples from adult patients with ND-AML (n=100) were analyzed by the qPCR rapid platform for stemness, cell state and target gene expression profiles. Subsets of these samples were used in the focused validation and response cohorts defined below.

#### HMA+Ven clinical response cohort

Samples were collected from ND-AML patients (n=58) treated with frontline HMA+Ven at OHSU with data for genomic, cytogenetic, and immunophenotypic characteristics and outcomes for response to HMA+Ven therapy. Survival data were available for all 58 patients in this cohort. Median time from AML diagnosis to commencing HMA+Ven induction was 8 days (interquartile range (IQR): 5-20). Median follow-up was 24.6 months from diagnosis. There were 38 deaths (66%) and median OS was 13.4 months (CI: 6.3-19.5). Clinical, genetic and immunophenotype data for all patients in this cohort are provided in Supplemental Table S1. Determinations of response and refractory states were obtained from patient records made by attending clinicians in line with NCCN guidelines.

#### Cell State assay validation cohort

Samples from adult AML patients (n=40) in the Beat AML study with RNAseq data were analyzed on both qPCR and research NanoString panels. Gene set enrichment scores from qPCR and NanoString data (see below) were compared to deconvoluted 30-gene cell-type-inferring scores (i.e., eigengenes) determined from RNAseq^12,19^. The patients in this cohort received 7+3 chemotherapy.

#### LSC17 assay validation

RNA samples (n=30) from ND-AML patients who received 7+3 chemotherapy and previously profiled for LSC17 signatures using a CLIA lab-developed NanoString assay as part of the reference cohort developed at PM were sent to OHSU for evaluation using qPCR technology. An analogous set of RNA samples (n=20) from ND-AML patients who received 7+3 chemotherapy at OHSU and previously characterized for LSC17 signatures using qPCR were sent to PM for measurement of LSC17 scores using the clinical NanoString assay. For qPCR-derived data, normalized gene expression was computed as delta Cq values for each gene compared to mean Cq of control genes, converted to fold-change (2^-ΔCq^), log2 transformed, multiplied by the previous reported regression coefficient weights for each gene^24^ and summed to calculate LSC17 gene scores for all 50 samples.

### TaqMan array card assay

Total RNAs (500 ng/sample, prepared as above) were reverse transcribed with Multiscribe (Thermo-Fisher). cDNAs were dispensed into customized TaqMan array cards containing pre-optimized primers, probes and reagents for amplification (Life Technologies) (Supplemental Table S2) and assayed in a QuantStudio 7 Flex (Applied Biosystems). The card layout, with at least two technical replicates for each gene, includes: 5 control genes, the top 25 ranked genes for each cell state signature^12^, and 10 apoptosis family genes. Data were quality processed through Design & Analysis v2.7 software (Applied Biosystems). Mean Cq values across technical replicates were compared to mean control gene Cq values (ΔCq). Stingscore values for enrichment of cell state signatures (as above) were calculated from ΔCq values for genes. Fold-changes in gene expression were calculated using the 2^-ΔΔCt^ formula. LinClass-7 score was calculated similarly to LSC17 scores as above using previously reported gene weights^20^. BCL2 gene expression ratios were calculated using the 2^-ΔCt^ formula. The BCL2 / (MCL1 + BCL2L*1*) ratio was calculated from ΔCq values as 2^(-1 ^*^ BCL2) / (2^(-1 ^*^ MCL1) + 2^(-1 ^*^ BCL2L1)).

### NanoString research assay

Total RNAs were extracted from freshly isolated bone marrow and peripheral blood mononuclear cells using the QIAGEN RNeasy Mini Kit with DNase treatment. NanoString cartridges were loaded with 125 ng of total RNA after probe hybridization with customized code sets prepared and optimized by NanoString for evaluation using the nCounter instrument. Code sets for this panel included seven control genes^10^ and the top 30 ranked genes from six cell state signatures^12^ (Supplemental Table S3). Raw data were processed using a pipeline of nCounter software and the nanostringr package^25^. Single-sample gene set enrichment analysis was performed using stingscores^26,27^ ranked with respect to stable control genes (*ACTB, GAPDH, VPS4A*).

### Ex vivo drug sensitivity assay

Testing for ex vivo drug sensitivity was performed as previously described ^22^. All ex vivo drug sensitivity tests were performed on freshly isolated mononuclear cells from patient samples collected at initial diagnosis. Viability assessments were made after 3 days of culture in the presence of drug.

### Statistical considerations

#### Correlation of gene signatures and drug sensitivity

Pearson correlation coefficients (“r”) and associated p-values were estimated (i) for each bulk RNAseq-derived cell type score (as computed in^19^) versus stingscores from qPCR or NanoString data and (ii) for LSC17 and LinClass-7 scores versus AUC values depicting sample sensitivity to Aza+Ven ex vivo. LSC17, LinClass-7 and BCL2 gene ratios were dichotomized according to the median value for comparison of Aza+Ven ex vivo AUC using a Wilcoxon rank sum test with Benjamini-Hochberg step-up correction for multiple comparisons. LSC17 score was compared based on achievement of CR on 7+3 treatment using a Wilcoxon rank sum test.

#### Clinical, genetic and gene expression features as predictors of HMA+Ven outcomes

AML-specific clinical, genetic, cytogenetic, and flow cytometry-detected characteristics were manually curated from patient electronic medical records. Outcomes of interest were composite CR (cCR), defined as achievement of CR or CRi (CR with incomplete blood count recovery) in response to HMA+Ven, and OS. Median follow-up, measured from AML diagnosis, was estimated by the reverse Kaplan-Meier (KM) method. Odds ratios (ORs) and receiver operating characteristic (ROC) curves and areas underneath these (AUCs) were computed from logistic regression models of cCR. OS was measured from AML diagnosis, with initiation of HMA+Ven considered the left-truncation time (to account for patient variability in entry times into the risk set). The KM method produced median OS estimates (and log-log 95% confidence intervals [CIs]) for median-dichotomized “low” vs. “high” BCL2 and BCL2/(MCL1 + BCL2L1) expression groups, with log-rank tests comparing KM curves. Hazard ratios (HRs) with accompanying Wald test p-values as well as concordance (C) indices and 1-year OS ROC curve (fromnearest neighbor estimation) AUCs for quantifying OS discrimination were calculated from Cox proportional hazards (PH) regression models^28^. The PH assumption was checked by visual examination of scaled Schoenfeld residuals. For both outcomes, a univariable model p-value threshold of 0.150 was used for consideration in multivariable (MVA) models, followed by application of Akaike Information Criteria (AIC) -based backward elimination separately for two feature sets: 1) clinical/genetic/immunophenotype variables assumed to be readily available to clinicians and 2) qPCR-derived gene expression variables from the Myelo-ID assay. The linearity assumption (checked with non-parametric smoothers such as LOESS or penalized splines), pairwise interaction terms (tested, separately, in the presence of the AIC-determined additive model), and multicollinearity (checked with Variance Inflation Factors) were assessed and considered satisfactory for the final MVA models. Statistical analyses were performed using R version 4.3.2 (R Core Team 2023).

## Results

### Rapid profiling of AML stemness, cell state and target expression

To develop a rapid method for determining tumor stemness and differentiation states using gene expression signatures, we evaluated a qPCR format with a customized panel containing pre-optimized primers (TaqMan Array cards). We customized the format to contain manufacturer-optimized gene sets that collectively measured signatures for leukemia stem cells (LSC17 ^10^); 6 AML cell states ^12^; and 10 genes encoding apoptosis proteins. The platform produces rapid, report-ready results within 48 hours of sample receipt, which is compatible with timelines for clinical decision making (Fig. 1A). We evaluated RNAs from 140 AML samples of which 100 were ND-AML patients on this platform (Supplemental Table S4). Representative samples for 6 of these AML patients reveal expression patterns for the LSC17 genes, cell state signatures and expression levels for apoptosis genes. Most samples showed prominent elevation of just one signature with some samples also exhibiting less pronounced elevation of an additional signature. In contrast, RNA isolated from a healthy donor bone marrow sample showed balanced expression of LSC17 and all six cell state signatures indicative of a normal functioning and heterogeneous blood compartment (Fig. 1B). Tumor cell state profiles with increased expression (as measured by z-score) of primitive gene sets distinguished profiles from those with differentiated cell states. For example, patient sample 17-01021 has an elevated LSC17 and HSC-like cell state profile; in contrast, patient sample 17-01083 has a high monocyte-like cell state profile. Across the complement of ND-AML patient samples that were also tested for ex vivo drug sensitivity (n=53), increased sensitivity to Aza+Ven aligned with more primitive tumor profiles is (Supplemental Figure S1).

**Figure 1.**
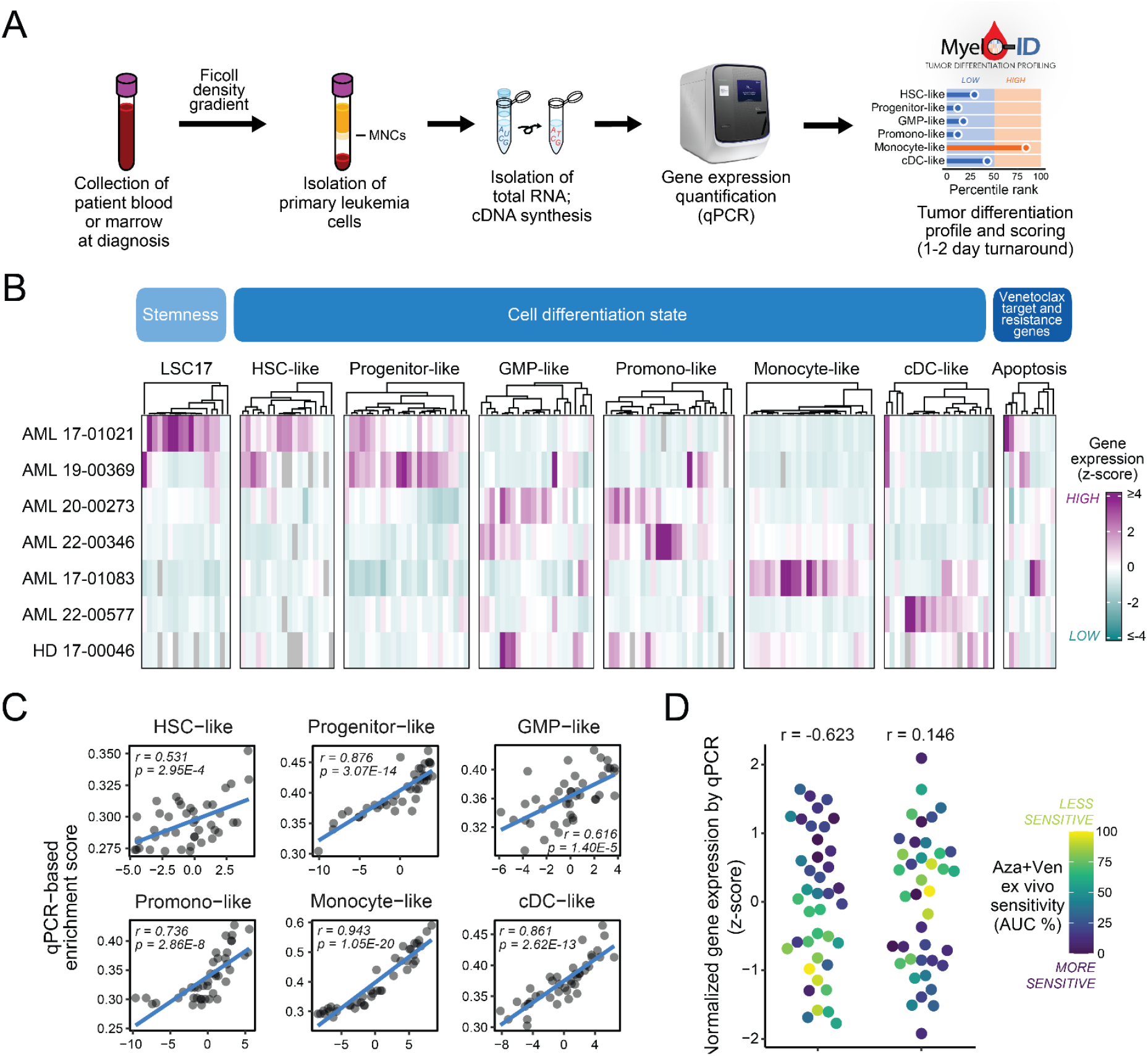
Rapid expression profiling defines AML stemness, cell state and target expression. **(A)** Workflow of gene expression profiling from sample acquisition to data output. **(B)** Heatmap of cell states identified by gene expression profiling for LSC17, cell state and apoptosis gene expression levels on 6 AML patient samples representative of each cell state and a healthy donor sample. Increased (orange) or decreased (blue) expression reflects z-scores standardized within each sample. Grey shading denotes undetectable level of gene expression. **(C)** Pearson correlation plots comparing cell state signatures derived by qPCR with bulk RNAseq eigengenes for AML samples (n=49). **(D)** Comparison of BCL2 and MCL1 levels determined by qPCR and correlated with ex vivo sensitivity to the combination of azacitidine plus venetoclax (Aza+Ven; n=53). Area under the dose response curve (AUC) was derived from a probit regression fit to a 7-point drug concentration series with cell viability measured by MTS-based assay after 3 days in culture. Values range from high (purple) to low (yellow) based on percentage of maximum AUC.

To examine the concordance between the data from this qPCR platform and RNA sequencing for the same 6 cell state signatures, we computed enrichment scores for these signatures using the stingscore method^26,27^ for the qPCR data and the previously published eigengene method ^19^ for the RNAseq data. Comparing these enrichment scores for 40 patient samples for which we have matched RNAseq and qPCR data revealed highly concordant values derived for the same patients for all 6 cell state signatures. Pearson r values range from 0.531 to 0.943 (p values range from 2.95e-4 to 1.05e-20; Fig. 1C). These data support the capability of gene expression profiling by qPCR to provide rapid quantitative assessments of tumor stemness and differentiation states concordant with those inferred from deconvolution of bulk RNA sequence data.

Prior studies have indicated a positive correlation between primitive cell states and ex vivo sensitivity to venetoclax^19,20^ or to Aza+Ven^22^. Moreover, the observation that expression levels for BCL2, the target of venetoclax, are higher in primitive cell states^15^ prompted the addition of genes for 10 apoptosis family members to the qPCR panel. Assessment of expression levels of BCL2 and MCL1, determined from qPCR data, demonstrated that elevated BCL2 levels and lower MCL1 levels were associated with ex vivo sensitivity to Aza+Ven as denoted by area under the dose response curve (AUC). (Pearson r: -0.623 and 0.146, respectively; Fig. 1D).

### Gene expression profiling adds predictive value to genetic features for HMA+Ven clinical outcomes

To determine the contribution of gene expression-based scores to patient response and survival on HMA+Ven therapy, we analyzed a 58-patient cohort of newly diagnosed AML patients treated with HMA+Ven from whom a diagnostic sample was available to evaluate by qPCR. Of these 58 patients, 56 were evaluable for clinical response; 37 (66.1%) attained a cCR. Among a panel of 108 clinical, genetic, and gene expression features (Fig. 2A and Supplemental Table S5), mutated *DNMT3A* (OR: 7.22 (95% CI: 1.74-49.80), p=0.016) was significantly associated with achieving cCR. In contrast, leukocytosis (OR: 0.16 (CI: 0.04 - 0.54), p=0.005) and high enrichment score for the promonocyte-like gene signature (OR: 0.19 (CI: 0.05 - 0.63), p=0.009) were associated with reduced likelihood of response (Fig. 2B). Among the flow cytometry assessments for cell differentiation ^6^, myeloid maturation markers (CD11b, CD15, CD64, CD65) trended towards an association with cCR although this was not statistically significant.

**Figure 2.**
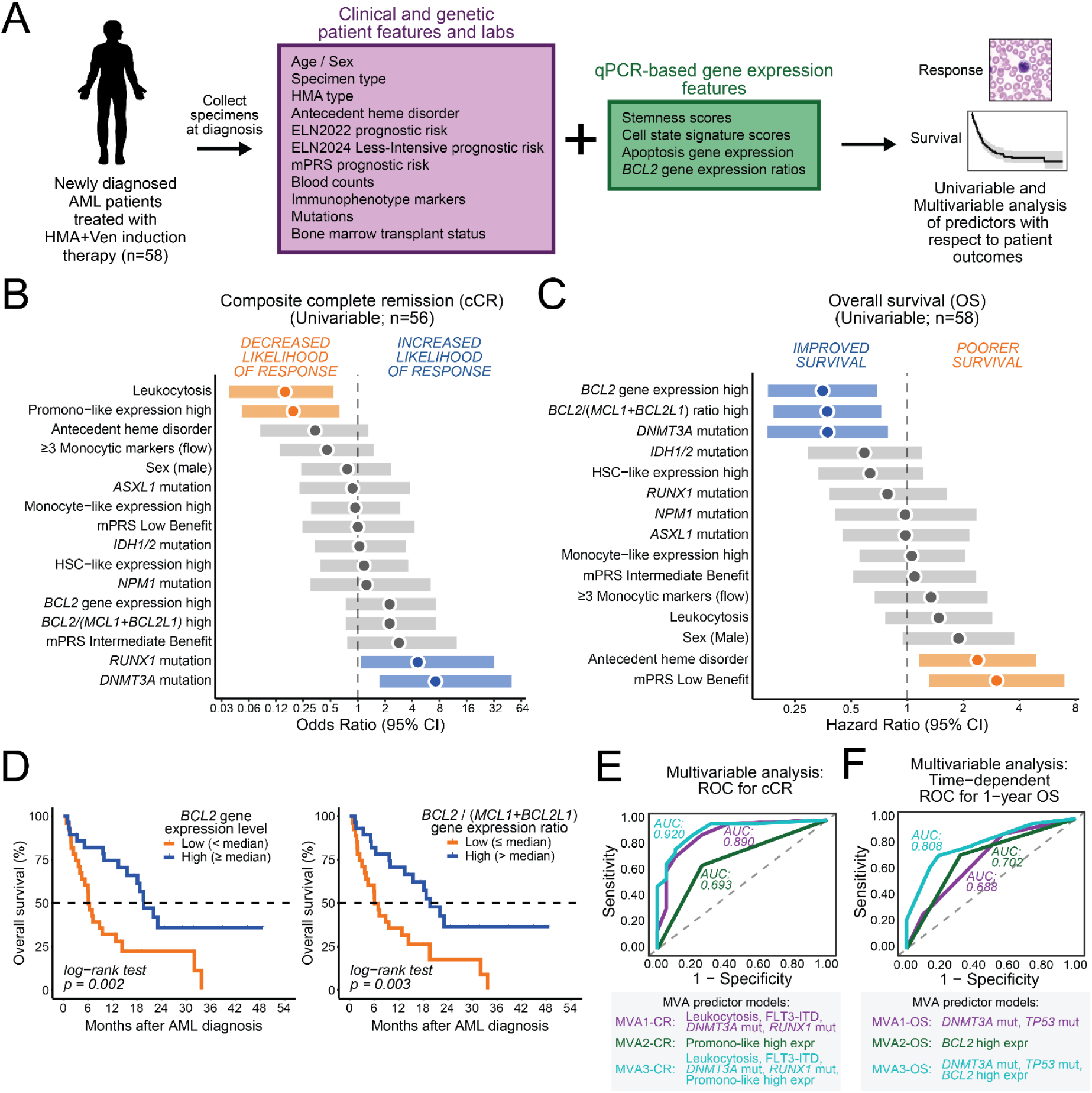
Gene expression profiling adds predictive value for HMA+Ven treatment outcomes. **(A)** Panel of 108 clinical, genetic, and gene expression features included as predictor variables in regression analyses with patient outcomes. (**B**) Forest plot depicting univariable logistic regression estimates for clinical, genetic, and gene expression features of ND-AML patients (n=56 evaluable) for achievement of composite complete remission (cCR). (**C**) Forest plot depicting univariable Cox regression estimates for clinical, genetic, and gene expression features for overall survival (OS) of ND-AML (n=58 evaluable). (**D**) Kaplan-Meier analysis for OS in patients with respect to BCL2 expression and to BCL2 / (MCL1+BCL2L1) gene expression ratio. (**E**) ROC curves from multivariable logistic regression models of cCR including AIC-driven backward selection-identified clinical, mutational, or immunophenotypic predictors (MVA1-CR; purple line); selected qPCR-based gene expression predictors (MVA2-CR; green line); and the combination of both sets pf predictors (MVA3 -CR; blue line). (**F**) One-year OS ROC curves from multivariable Cox models including clinical, mutational, or immunophenotypic predictors (MVA1-OS, purple line); selected qPCR-based gene expression predictors (MVA2-OS, green line); and when combining both sets of predictors chosen from the AIC-based model selection strategy (MVA3-OS, blue line).

With respect to overall survival (OS), mutated *DNMT3A* was associated with improved OS (median: 21.9 vs 7.2 months; HR: 0.37 (CI: 0.18-0.79), p=0.010), whereas antecedent hematologic disorder (5.9 vs 15.4 months; HR: 2.38 (CI: 1.16-4.90), p=0.019) was associated with shorter OS (Fig. 2C and Supplemental Table S5). The association of mutated *DNMT3A* with OS is concordant with findings from two clinical cohorts on venetoclax-based therapies ^5,29^. mPRS risk classification showed median OS of 19.6, 14.3, and 6.0 months for high, intermediate, and low benefit groups, respectively. Compared to high benefit patients, those with mPRS low benefit classification (i.e., *TP53*-mutated) had significantly inferior OS (HR: 3.02 (CI: 1.31-6.96), p=0.010). Gene expression scores also showed predictive value for OS; high BCL2 expression was the most significant predictor of survival among all features examined (HR: 0.35 (CI: 0.18-0.69), p=0.002). Furthermore, increased relative expression of BCL2, as measured using the gene expression ratios for BCL2 / BCL2L1 (HR: 0.48 (CI: 0.25-0.93), p=0.028) and BCL2 / (MCL1+BCL2L1) (HR: 0.37 (CI: 0.19-0.73), p=0.004), was also significantly associated with improved OS. Kaplan-Meier analysis showed significantly longer survival in patients with high BCL2 expression or high BCL2 / (MCL1+BCL2L1) gene expression ratio (median OS in both cases: 19.6 vs 6.0 months) (Fig. 2D). Thus, RNA levels of BCL2, the target of venetoclax, is a predominant indicator of OS for HMA+Ven treated patients.

We used a series of AIC-selected multivariable models to comparatively evaluate the contributions of clinical, genetic, and gene expression features for predicting cCR and OS. Model 1 included multivariable model-selected clinical and genetic variables. cCR Model 1 included leukocytosis (OR: 0.07, p=0.004), *FLT3-ITD* (OR: 12.5, p=0.068), mutated *DNMT3A* (OR: 17.1, p=0.007), and mutated *RUNX1* (OR: 12.6, p=0.018) and had an AUC on a Receiver Operating Characteristic (ROC) curve of 0.890. Model 2 represented the multivariable model-selected gene expression features. For cCR, Model 2 included the promonocyte-like gene score alone and demonstrated an AUC of 0.693. Inclusion of all features from Models 1 and 2 (Model 3) revealed that combining clinical, genetic, and gene expression features improved the ability to discriminate and predict cCR (ROC AUC: 0.920; Fig. 2E and Supplemental Table S6). Using a similar approach to quantify and compare clinical, genetic, and gene expression features for predicting OS, consideration of only clinical and genetic variables led to an MVA Model 1 comprising mutated *DNMT3A* (HR: 0.42, p=0.025) and mutated *TP53* (HR: 2.43, p=0.020) with a C-index of 0.645 and 1-year ROC AUC of 0.688. The gene expression MVA Model 2 involved only BCL2 gene expression and showed a C-index of 0.630 and 1-year ROC AUC of 0.702. Inclusion of high BCL2 expression with Model 1 features (Model 3) (HR: 0.32, p=0.001) enhanced OS discrimination, yielding a C-index of 0.713 and 1-year ROC AUC of 0.808 (Fig. 2F and Supplemental Table S6). Hence, the multivariable models indicate that cell state scores and BCL2 expression alone predict clinical outcomes in a comparable fashion to the most predictive clinical and genetic features, and inclusion of these gene expression levels with clinical and genetic features adds prognostic value.

### Illustration of gene expression profiling value in an individual case

To demonstrate the utility of cell state signatures and BCL2 expression levels for indicating responsiveness to HMA+Ven therapy, we profiled the cell state of a diagnostic sample from a newly diagnosed AML patient (6705). For this patient, the initiation of HMA+Ven produced a modest reduction in bone marrow blasts that was refractory after 2 cycles of treatment (Fig. 3A). The clinical and genetic features at diagnosis indicated High Benefit by mPRS criteria suggesting the patient was likely to respond to HMA+Ven treatment (Fig. 3B). However, gene expression profiling with Myelo-ID showed that the patient had a differentiated cell state profile (high promonocyte-like, monocyte-like and cDC-like scores, Fig. 3C) and low expression levels of BCL2 (Fig. 3D). For patient 6705, gene expression analysis was concordant with the observed insufficient clinical responsiveness to HMA+Ven therapy. In contrast, for patients classified as mPRS Low Benefit, the strong independent prognostic value of high BCL2 gene expression at diagnosis identified in multivariable analysis suggests an improved ability to stratif y outcomes.

**Figure 3.**
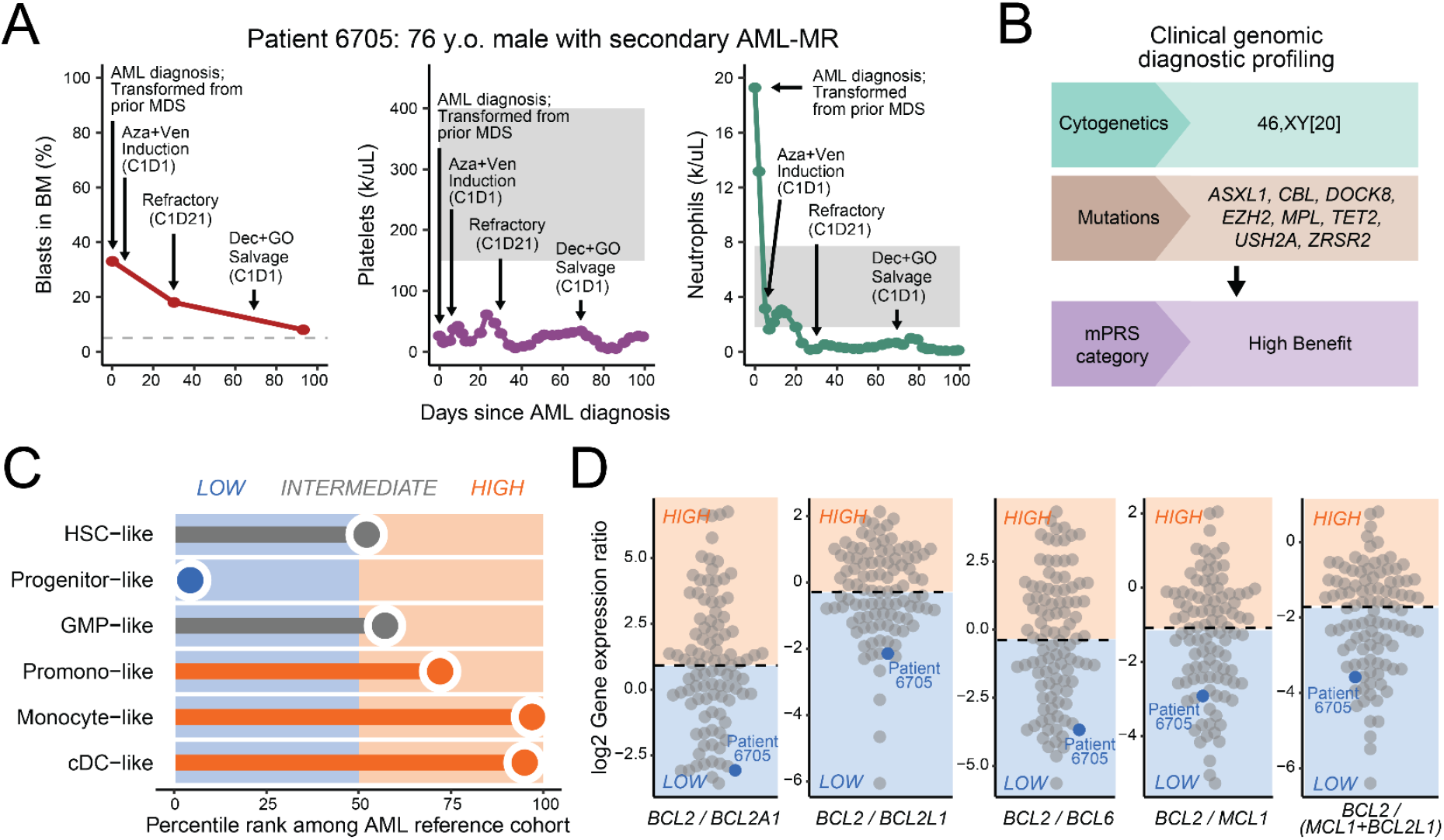
Cell state profiling adds context for HMA+Ven therapy selection. **(A)** Bone marrow blast % and platelet counts for ND-AML patient 6705 following HMA+Ven treatment. **(B)** Clinical genomic profile for patient 6705 at diagnosis indicate categorization as mPRS High Benefit. **(C)** qPCR-based cell state profile for patient 6705 at diagnosis. **(D)** Ratios of expression levels for BCL2 relative to BCL2A1, BCL2L1, BCL6, MCL1, and MCL1+BCL2L1 at diagnosis.

### Gene expression associations with ex vivo HMA+Ven sensitivity and immunophenotypic features

Given that BCL2 expression was such a strong predictor of HMA+Ven sensitivity in the clinical outcome analysis, we compared Myelo-ID results with ex vivo Aza+Ven sensitivity. In a cohort of newly diagnosed AML patient samples (n=53), we found an association of increased ex vivo Aza+Ven sensitivity that aligned with elevated gene expression levels for BCL2 as well as anti-apoptotic genes controlled by the TP53 pathway. Ex vivo HMA+Ven insensitivity aligned with genes that contribute to mitochondrial resistance (e.g., MCL1, BCL2A1^15^) or transcriptional repression (e.g. BCL6^30^) (Fig. 4A). High ratios of BCL2 expression compared to expression of other apoptosis genes were significantly associated with ex vivo HMA+Ven sensitivity (Fig. 4B), which is consistent with the association of clinical responsiveness to Ven with target protein ratios by flow cytometry^31^ and prospective ex vivo screening in the VenEx trial (NCT04267081)^32^. Myelo-ID profiling also provides nuanced context to augment immunophenotype results generated by flow cytometry. Sensitivities of HMA+Ven ex vivo among patient samples with respect to primitive (CD117) and monocytic (CD11b) markers show a broad range of effectiveness (Fig. 4C). Further stratification of HMA+Ven sensitivity ex vivo in CD117+ or CD11b+ patients by BCL2 and cell state enrichment scores results in enhanced delineation of responsiveness compared with the overlapping ranges derived from flow cytometry alone (Fig. 4D). These findings underscore the ability of cell state profiling to augment interpretationof clinical flow cytometry datafor predicting responsiveness to HMA+Ven treatment.

**Figure 4.**
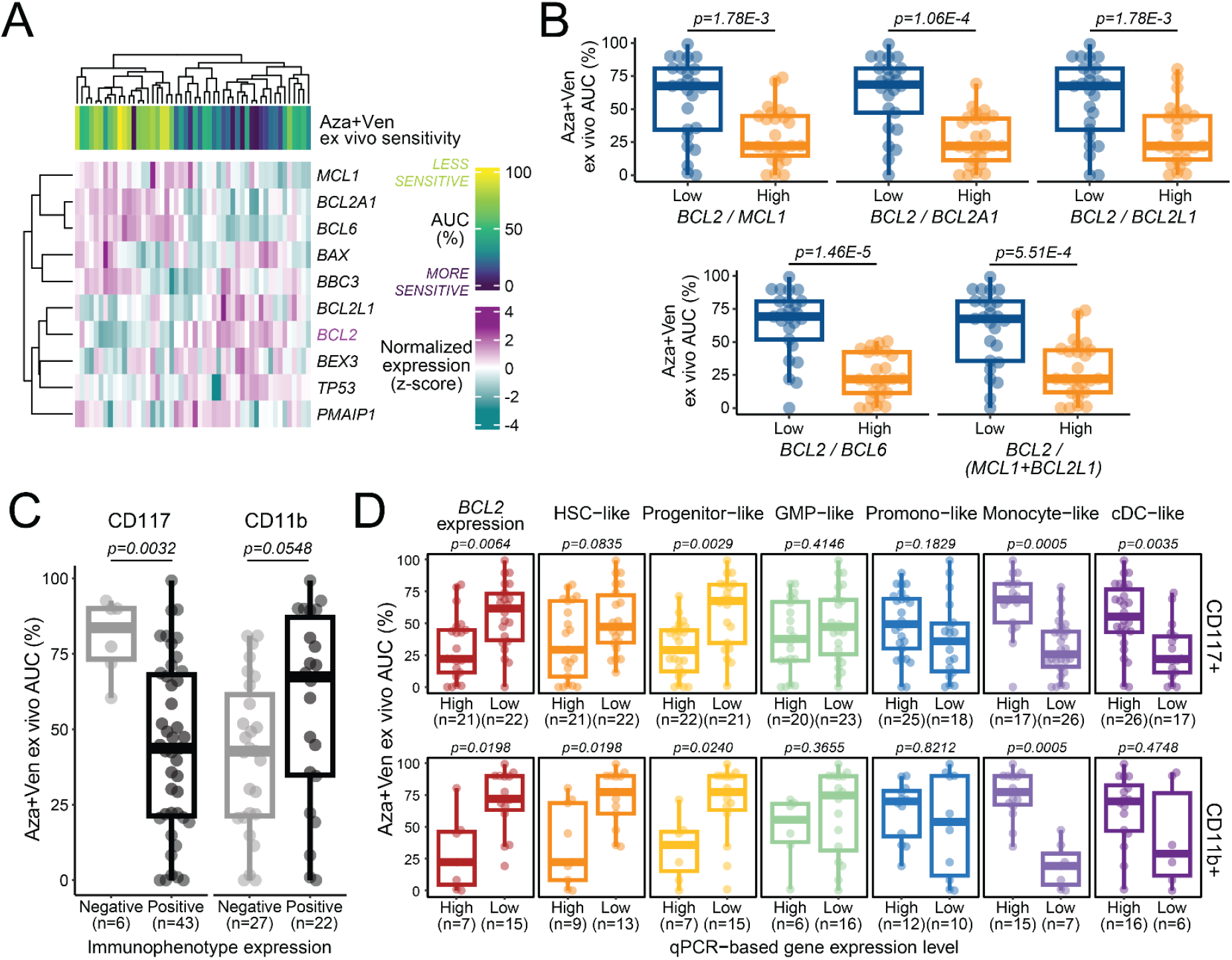
qPCR-derived *gene* expression scores associate with ex vivo sensitivity to HMA+Ven and immunophenotype features. **(A)** Heatmap depicting expression levels for 10 apoptosis genes aligned with ex vivo sensitivity to Aza+Ven (n=53). **(B)** Ratios of expression levels for BCL2 relative to MCL1, BCL2A1, BCL2L1, BCL6 and MCL1+ BCL2L1 plotted with respect to Aza+Ven sensitivity ex vivo. The median value for each gene expression ratio was used as a cut point for High vs Low determination. AUC % denotes area under the dose response curve as a percentage of maximum AUC. (**C**) Aza+Ven sensitivity ex vivo among patient samples with respect to primitive (CD117) and monocytic (CD11b) markers (n=49 evaluable). (**D**) Aza+Ven sensitivity ex vivo in patients positive for CD117 or CD11b according to relative enrichment scores for BCL2 and 6 cell states. Expression scores are binned as median-dichotomized values for consistency with the clinical response cohort in Fig. 2.

### Expression profiling for LSC17 expands the utility of the rapid platform

We evaluated the utility of gene expression profiling with the stemness component of Myelo-ID for prognosticating responsiveness to intensive chemotherapy in AML. To compare LSC17 scores as obtained by qPCR with respect to those by a clinically approved NanoString assay conducted at Princess Margaret (PM) Cancer Center, we carried out a reciprocal exchange of samples from AML patients treated with intensive chemotherapy. Twenty samples from OHSU, where LSC17 scores were initially evaluated using qPCR technology (n=20 samples) were sent to PM for Nanostring analysis, and thirty samples from the PM AML reference cohort^10^ were assessed by qPCR at OHSU. Scores measured using the two methods were highly concordant (Pearson r=0.9313; p<2.2e-16; Fig. 5A and Supplemental Fig. S2). A similar comparison was performed for cell state scores as measured by qPCR or Nanostring which indicated high concordance and stability of expression across batch runs (Supplemental Fig. S3). Thus, either platform provides a rapid and robust means to determine gene expression profiles.

**Figure 5.**
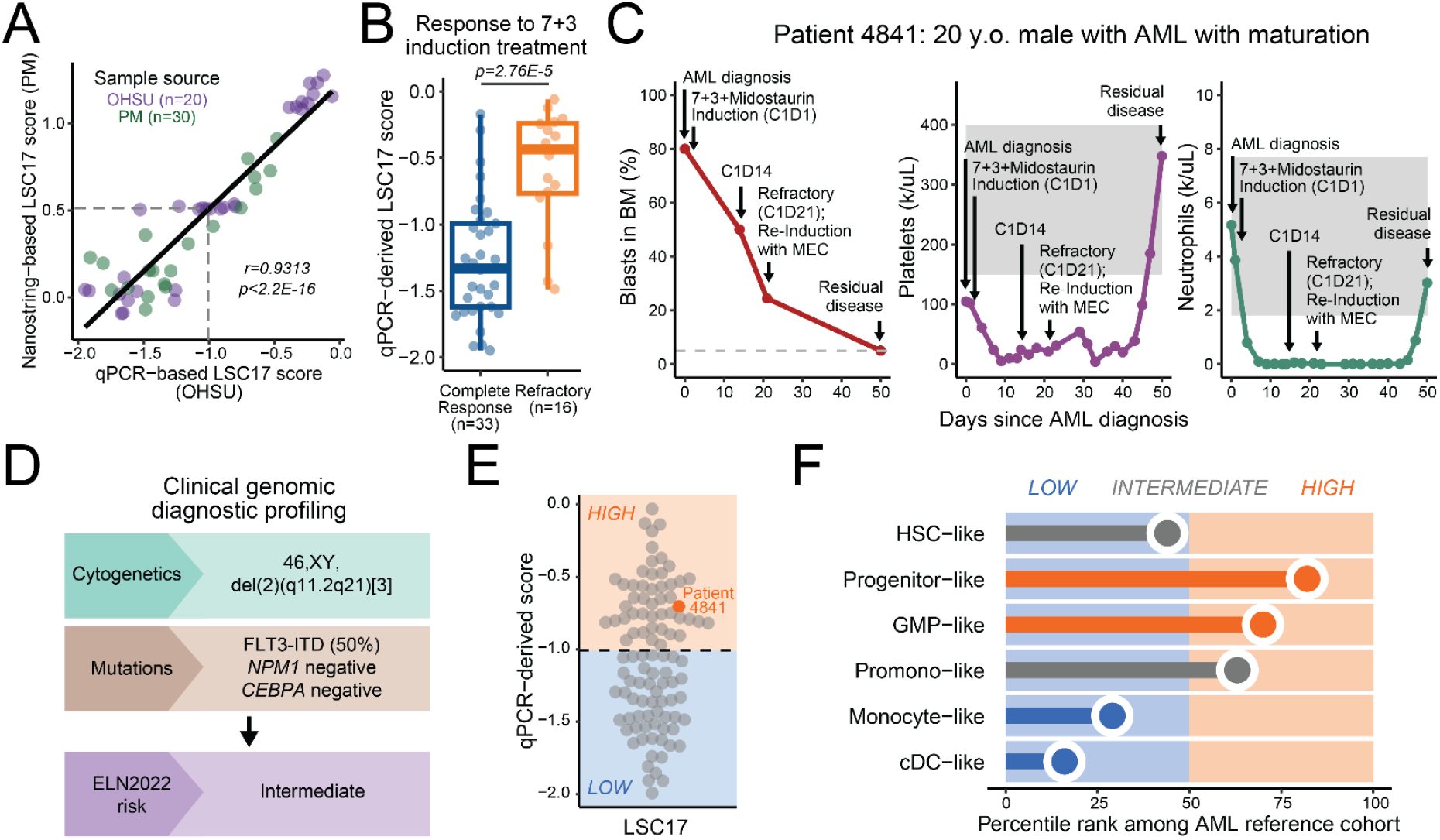
qPCR profiling for LSC17 and cell state augments clinical features. **(A)** Pearson correlation plot of LSC17 scores determined by qPCR versus Nanostring for 50 patient samples: 30 samples from the LSC17 validation cohort assembled at Princess Margaret Hospital (PM; purple) and 20 AML patient samples from OHSU (green) were evaluated at both institutions using NanoString technology at PM and qPCR technology at OHSU. Dashed lines denote the median (used as a cut point for Low vs High group determination) scores determined by each method. **(B)** Comparison of qPCR derived scores for LSC17 plotted for patients achieving a CR on 7+3 induction therapy (n=33) versus LSC17 scores for patients refractory to therapy (n=16). **(C)** Blood compartment counts for bone marrow blast %, platelets and neutrophils plotted over two cycles of 7+3 therapy for patient 4841. Following the failure of standard induction chemotherapy, the patient was re-induced with mitoxantrone, etoposide, and intermediate dose cytarabine (MEC) therapy. **(D)** Clinical genomic profile for patient 4841 at diagnosis. (**E)** qPCR-derived LSC17 score for patient 4841 at diagnosis (highlighted in orange) plotted with the LSC17 scores for 100 ND-AML samples. **(F)** Cell state profile of patient 4841 at diagnosis.

Assessment of cumulative patient responses to 7+3 chemotherapy in this 50-patient cohort indicated that lower LSC17 scores were associated with CR on therapy, a result consistent with prior findings^24^ (Fig. 5B; median of -1.33 vs -0.435; p=2.76E-5). As a use case illustration, measurement of LSC17 score and cell state determination was performed by qPCR on a diagnosis sample from an AML patient before treatment with 7+3 chemotherapy and midostaurin. Over two cycles of treatment, bone marrow blasts in AML patient 4841 were reduced from 80% to only 25% (Fig. 5C). At this time, the clinician notes indicated a change to salvage therapy due to refractory disease. At diagnosis, the patient was classified as intermediate risk (ELN 2022) with a FLT3-ITD mutation (Fig. 5D). However gene expression profiling revealed a high LSC17 score and a primitive cell state profile, both associated with non-responsiveness to 7+3 therapy (Fig. 5E, F), illustrating the added value of gene expression profiling for guiding therapy selection.

Although the LSC17 stemness signature has prognostic value in the setting of intensive chemotherapy, there was no significant association between LSC17 scores and HMA+Ven response either clinically or using ex vivo assessments (Fig. 5 and Supplemental Fig. S4A). However, LinClass-7, a subset of genes derived from LSC17^20^, was strongly associated with ex vivo sensitivity to HMA+Ven, with high score samples showing greater sensitivity (Supplemental Fig. S4B). These findings are concordant with prior findings for venetoclax using deconvoluted RNAseq data^19,20^ and indicate an additional gene expression feature to aid with predicting HMA+Ven responsiveness.

## Discussion

Gene expression profiling is in wide use for cancers such as breast ^33^ and prostate^34^, enabled by the incidence of these malignancies to power statistically-rigorous predictors of progression and survival. This paradigm has yet to be similarly realized for AML, a rarer cancer with heterogenous genetic origins. The LSC17 score, derived from functionally validated leukemic stem cell populations, is associated with survival following standard (7+3) chemotherapy and has been developed into a clinically available NanoString-based diagnostic assay^24^. Despite the further validation of the prognostic value of LSC17 in a prospective trial^35^, this score has not been widely implemented in diagnostic evaluations. The inclusion of LSC17 genes with key genes that predict response to HMA+Ven (apoptosis, AML tumor cell state) into one qPCR panel extends the application to a broad spectrum of AML patients treated with the two primary standard-of-care regimens through a comprehensive view of stemness, cell state, and drug target expression (Fig. 6).

**Figure 6.**
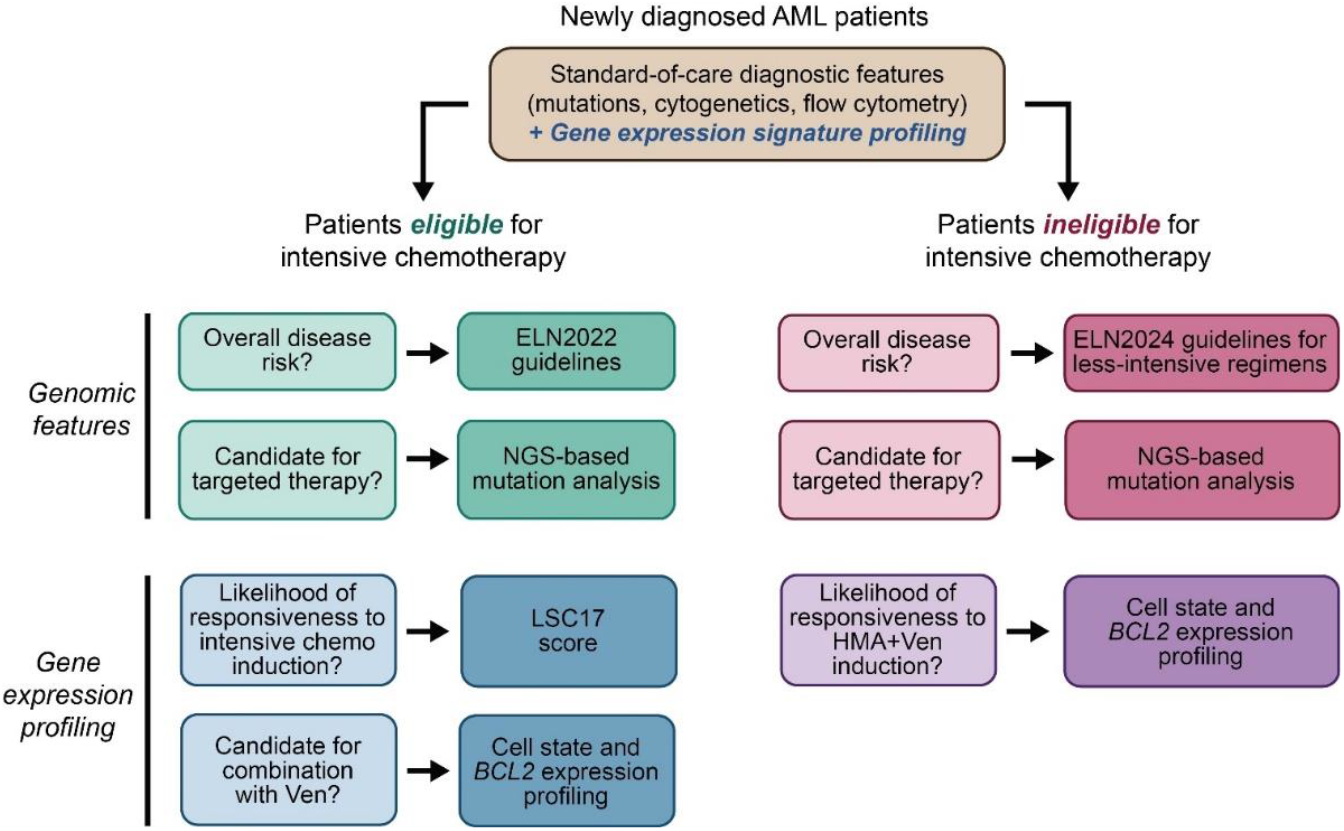
Implementation scheme for gene expression profiling in therapy selection. Representation of AML therapy selection based on patient fitness for intensive chemotherapy. In instances where NGS-based detection reveals mutations amenable to targeted drugs, incorporation of gilteritinib (*FLT3*), ivosidenib (*IDH1*), enasidenib (*IDH2*), or revumenib (*KMT2A*-rearranged) may offer improved benefit. ELN2022 guidelines^6^; ELN2024 Less-intensive guidelines^8^.

Prior studies using BH3 profiling^36,37^ or flow cytometry^31^ have correlated BCL2 family protein levels with HMA+Ven response and event-free survival. A key finding from this study is the demonstration that BCL2 transcript levels from bulk mononuclear cells at diagnosis predicts OS for HMA+Ven treatment. Patient survival was also stratified when evaluating BCL2 gene expression relative to MCL1 and BCL2L1, however, this did not improve the predictive power obtained from measuring *BCL2* transcript alone. Given the increasing use of venetoclax-based therapies for AML therapy in both adult and pediatric care, this simple detection method that requires only bulk RNA from diagnostic tumor tissue may facilitate increasingly rational therapeutic use of venetoclax through distributed application of this highly feasible diagnostic.

Gene expression profiling of AML tumors using Myelo-ID offers a standardization not captured by current flow cytometry methods, owing to the larger and more diverse collection of genes that can be assessed at the transcript level as well as more quantitative representation of the data. Correlation of LSC17 scores determined by qPCR with those obtained by the NanoString assay developed at PM establishes the utility of qPCR for measuring LSC17 and LinClass-7 scores. The value of standardizing results of gene expression-based laboratory tests is critical to consistent interpretation and broader implementation, as exemplified in the established guidelines and international standards for BCR-ABL1 transcript level testing^38^. To facilitate standardization and implementation broadly, we used commercial reagents, a widely used qPCR platform, and a similar high/low expression level cut point to that developed for LSC17. BCL2 expression assessment is similar to other gene expression-based tests that are in clinical workflows as validated and useful biomarkers. Gene expression adds value as a standardized measure of target expression. Flow cytometry, as used clinically in AML diagnosis, provides immunophenotype characterization as interpreter-dependent binary or categorical views of cell surface marker presence, but is never provided as a quantitative metric. Immunophenotyping is not included as a risk stratification feature in ELN guidelines for either standard 7+3 or HMA+Ven induction therapy. ELN guideline-defined sets of immunophenotypic markers were included in our univariable analysis of HMA+Ven treated patients but were not significant predictors of response or survival. However, a retrospective study using flow cytometry data that does utilize mean fluorescence intensities as continuous variables has shown predictive value of flow data used in this way (Lachowiez et al., 2025, *Blood Cancer Discovery*, in press).

For AML patients treated with HMA+Ven, however, there is currently no validated gene expression-based biomarker predictive of therapy response. Both qPCR- and NanoString-derived tumor cell state enrichment correlated with RNAseq methods^19,20^. Among cell state signatures assessed here, only the promonocyte-like gene set was significantly associated with decreased likelihood of achieving cCR on HMA+Ven. While this finding illustrates the value for gene expression signatures in predicting clinical outcome, larger and validation cohorts will be needed to further delineate the cell states associated with outcomes to HMA+Ven or other therapeutic regimens. Given the heterogeneity of AML tumors, Myelo-ID adds context to the interpretation of an individual tumor. In some cases, patients express more than one cell state signature at high levels, which may indicate multiple clones in the tumor. Reduced levels of BCL2 have been associated with AML tumor differentiation and reduced sensitivity to venetoclax in prior studies ^15-18,23^. Expansion of resistant subclones with mutations are also observed at the time of resistance^21,39^.Recent descriptions of inflammatory gene signatures^40,41^ and inflammatory memory stem cells^42^ are also aspects of tumor complexity that may help discern responsiveness in these complex profile scenarios. Development of a larger cohort will enable prospective testing and assay refinement.

Our findings demonstrate that Myelo-ID profiling enhances molecular feature-based prediction of AML patient outcomes in the context of existing diagnostic tests. The advantages of qPCR array card technology for gene expression profiling include its relative widespread use, suitability for single-sample runs, and short turn-around times (24-48 hours) compatible with clinical decision making. Integration of gene expression profiles with clinically prognostic mutations may better inform treatment selection and improve outcomes in a manner similar to the International Prognostic Scoring System for Myelodysplastic Syndromes (IPSS-M) test which integrates hematologic parameters, cytogenetic abnormalities and somatic mutations for 31 genes into a patient -specific risk score with clinical utility^43^. Rapid AML stratification based on gene expression classifications permits the up-front identification of patients who are most likely to benefit from standard of care therapies or other novel Ven-inclusive treatment regimens. For a cancer as heterogenous as AML, as diagnostic tools and the therapeutic landscape evolve, the ability to use all available clinical and genetic patient features to guide treatment selection may enable improved depth and duration of response in patients.

## ACKNOWLEDGEMENTS

The authors thank the patients treated at Oregon Health & Science University and Princess Margaret Cancer Centre for the generous use of their tissue samples. This work was supported by National Institutes of Health (NIH), National Cancer Institute (NCI) grant U54CA224019, NCI award R01CA262758 (J.W.T., S.E.K.), NCI Cancer Center Support Grant P30CA069533 (S.E.K), Oregon Clinical and Translational Institute (S.E.K), the Waldman Family Fund for AML Research (J.W.T.), the George Ettelson Endowed Professorship In Acute Myeloid Leukemia Research (J.W.T.), the Mark Foundation for Cancer Research (J.W.T.), and the Silver Family Foundation (J.W.T.).

## Data Sharing Statement

Information and requests for resources and reagents should be directed to Dr. Jeffrey Tyner. RNAseq data for patient samples are available at dbGaP under study ID 30641.

## Authorship Contributions

Study Design: CAE, SEK, SA, JED, JWT; Assay implementation: SEK, CAE, AM, NL, CL; Data Analysis: CAE, SA, AK, CL, DB, SKM, SWKN, JCYW; Manuscript drafting and editing SEK, CAE; AM, JCWY, SWKN, AK, CL, JWT.

## Supplemental Material

**Supplemental Figure S1. Heatmap of newly diagnosed AML patient profiles for stemness, cell state and apoptosis gene expression aligned with ex vivo Aza+Ven sensitivity**. Genes within each set are clustered by expression across all 53 AML samples and one healthy donor (17-00046). Normalized expression is shown as z-scores for each gene. Ex vivo sensitivity to Aza+Ven was determined from MTS-based viability assay on freshly isolated mononuclear cells. Area under the drug response curve (AUC) is shown as a percentage of the maximum possibility value for the surveyed dose range; lower AUC values indicate greater sensitivity. Grey shading denotes data not available.

**Supplemental Figure S2. Consistency between clinical Nanostring and qPCR-based LSC17 score gene expression and interpretation. (A)** Gene expression distributions for each of 50 primary AML patient samples profile using both the clinical Nanostring LSC17 assay and the qPCR TaqMan array card panel. Relative expression levels of genes in the LSC17 signature and the set of five control genes are shown in blue and gray, respectively, with boxplots highlighting the interquartile range. While control gene expression remained comparable among all samples, a range of expression levels was detected for LSC17 genes, prior to calculating weighted score. **(B)** Contingency table comparing binary calls for LSC17 score for matched samples evaluated by either the clinical Nanostring LSC17 assay or the qPCR TaqMan platform. Calls of “High” or “Low” correspond to values above or below the reference median score values of 0.51 and -1.01 for the clinical Nanostring and qPCR assays, respectively. Frequencies were compared by McNemar Chi-square test without Yates correction. **(C)** Contingency table comparing concordance between qPCR-derived LSC17 score call with clinical responsiveness to 7+3 induction treatment. Assay sensitivity and specificity are shown. True positive results were instances where qPCR-derived LSC17 score call was high and the patient was refractory to clinical treatment. Abbreviations: TP, true positive; FP, false positive; FN, false negative; and TN, true negative.

**Supplemental Figure S3. Consistency of research Nanostring- and qPCR-based gene expression distribution by sample and batch. (A)** Concordance of cell state stingscores with qPCR versus Nanostring platforms (n=40). Pearson r values are indicated. (**B**) Distribution of normalized gene counts for 40 AML patient samples profiled with the research Nanostring panel. Samples are grouped according to cartridge batch, with boxplots highlighting the interquartile range of all genes for a given sample. **(C)** Distribution of normalized gene expression for 140 AML patient samples profiled using the qPCR TaqMan array card assay (upper panel). Samples are grouped according to reagent batch order of TaqMan cards. Expression (mean Cq) values for each of five control genes on the qPCR panel are shown for each of the same samples as above, in the same order from left to right (lower panel).

**Supplemental Figure S4. Correlation of qPCR-based stemness scores with ex vivo Aza+Ven sensitivity**. **(A)** Comparison of qPCR-derived scores for LSC17 versus ex vivo sensitivity to HMA+Ven for ND-AML samples (n=100). The median cut point for High vs Low LSC17 scores was established in Fig. 1B. **(B)** Comparison of qPCR-derived scores for LinClass-7 versus sensitivity to ex vivo HMA+Ven for ND-AML samples (n=100). The median cut point for High vs Low LinClass-7 scores was determined using the median score, as in ^20^.

**Supplemental Table S1. Clinical annotations for OHSU newly diagnosed AML patients receiving induction with either intensive chemotherapy or with HMA+Ven**.

**Supplemental Table S2. TaqMan gene assay reagent list. Supplemental Table S3 Nanostring gene assay reagent list**.

**Supplemental Table S4. Gene expression scores for patients analyzed by data type and analysis cohort**.

**Supplemental Table 5. Univariable analysis of clinical, genetic, and gene expression features to predict achievement of cCR or OS in HMA+Ven-treated AML patients**.

**Supplemental Table 6. Multivariable analysis of clinical, genetic, and gene expression features to predict achievement of cCR or OS in HMA+Ven-treated AML patients**.

