## Supplemental Figures S1-S4 for "A Rapid Gene Expression Profiler Classifies AML Tumor Responsiveness to Standard Therapies"

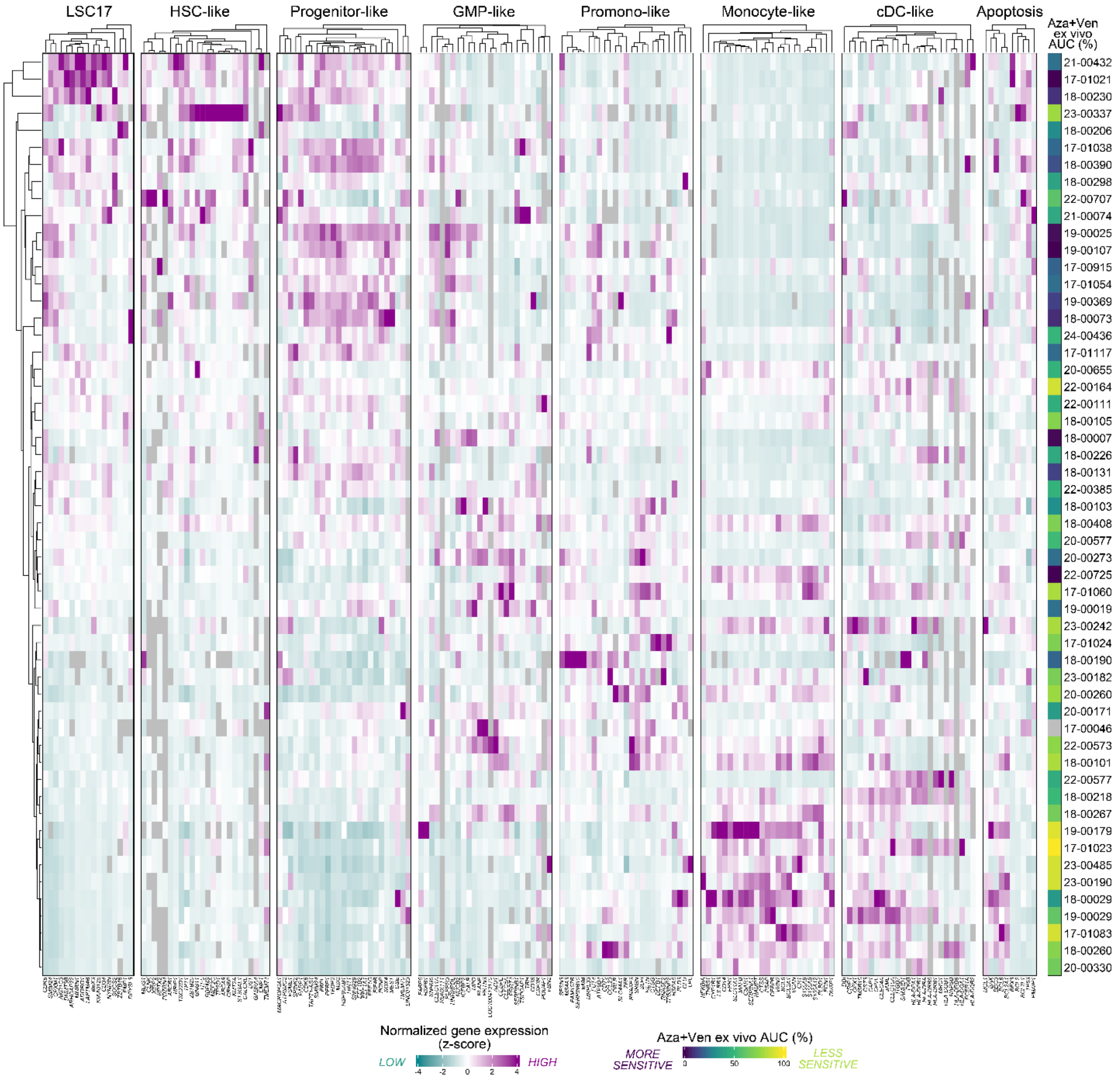


**Supplemental Figure S1. Heatmap of newly diagnosed AML patient profiles for stemness, cell state and apoptosis gene expression aligned with ex vivo Aza+Ven sensitivity.** Genes within each set are clustered by expression across all 53 AML samples and one healthy donor (17-00046). Normalized expression is shown as z-scores for each gene. Ex vivo sensitivity to Aza+Ven was determined from MTS-based viability assay on freshly isolated mononuclear cells. Area under the drug response curve (AUC) is shown as a percentage of the maximum possibility value for the surveyed dose range; lower AUC values indicate greater sensitivity. Grey shading denotes data not available.

A

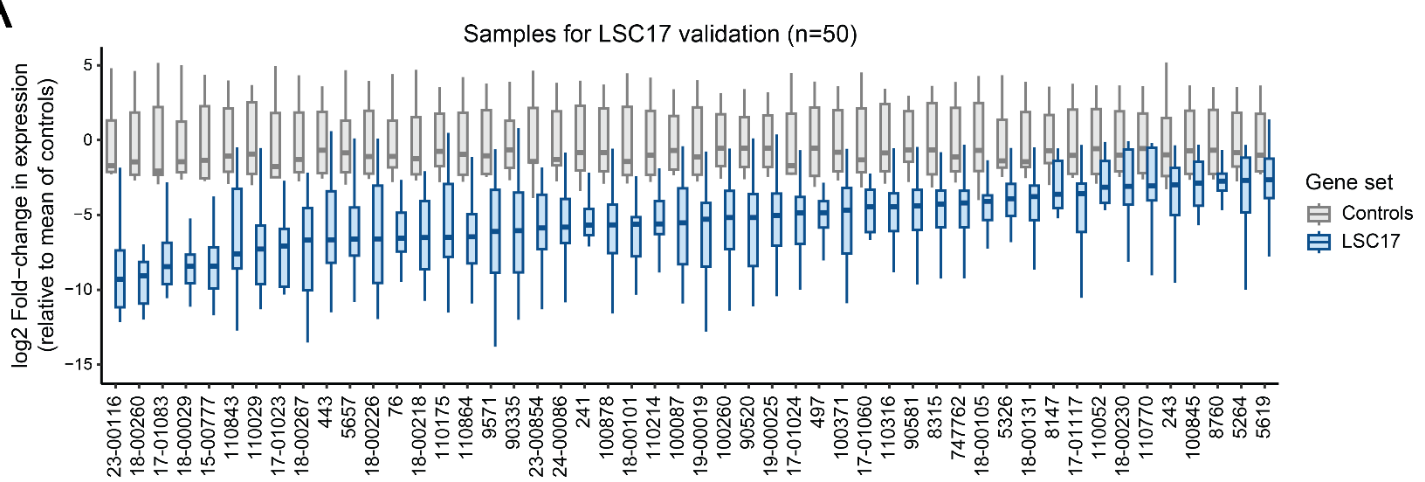


B

| | | Clinical Nanostring LSC17 assay: High | Clinical Nanostring LSC17 assay: Low |
|---|---|---|---|
| qPCR TaqMan LSC17 | High | 19 | 3 |
| | Low | 2 | 26 |

McNemar
Chi-square test
w/o Yates correction:
0.2
p=0.6547

Concordance rate:
45/50 (90%)

C

| | | Response to 7+3 induction: Complete response | Response to 7+3 induction: Refractory |
|---|---|---|---|
| qPCR TaqMan LSC17 | Low | 24 | 3 |
| | High | 9 | 13 |

Sensitivity
(TP / (TP + FN)):
0.812

Specificity
(TN / (FP + TN)):
0.727

**Supplemental Figure S2. Consistency between clinical Nanostring and qPCR-based LSC17 score gene expression and interpretation. (A)** Gene expression distributions for each of 50 primary AML patient samples profile using both the clinical Nanostring LSC17 assay and the qPCR TaqMan array card panel. Relative expression levels of genes in the LSC17 signature and the set of five control ge nes are shown in blue and gray, respectively, with boxplots highlighting the interquartile range. While control gene expression remained comparable among all samples, a range of expression levels was detected for LSC17 genes, prior to calculating weighted score. **(B)** Contingency table comparing binary calls for LSC17 score for matched samples evaluated by either the clinical Nanostring LSC17 assay or the qPCR TaqMan platform. Calls of “High” or “Low” correspond to values above or below the reference median score values of 0.51 and -1.01 for the clinical Nanostring and qPCR assays, respectively. Frequencies were compared by McNemar Chi-square test without Yates correction. **(C)** Contingency table comparing concordance between qPCR-derived LSC17 score call with clinical responsiveness to 7+3 induction treatment. Assay sensitivity and specificity are shown. True positive results were considered to be instances where qPCR-derived LSC17 score call as high and the patient was refractory to clinical treatment. Abbreviations: TP, true positive; FP, false positive; FN, false negative; and TN, true negative.

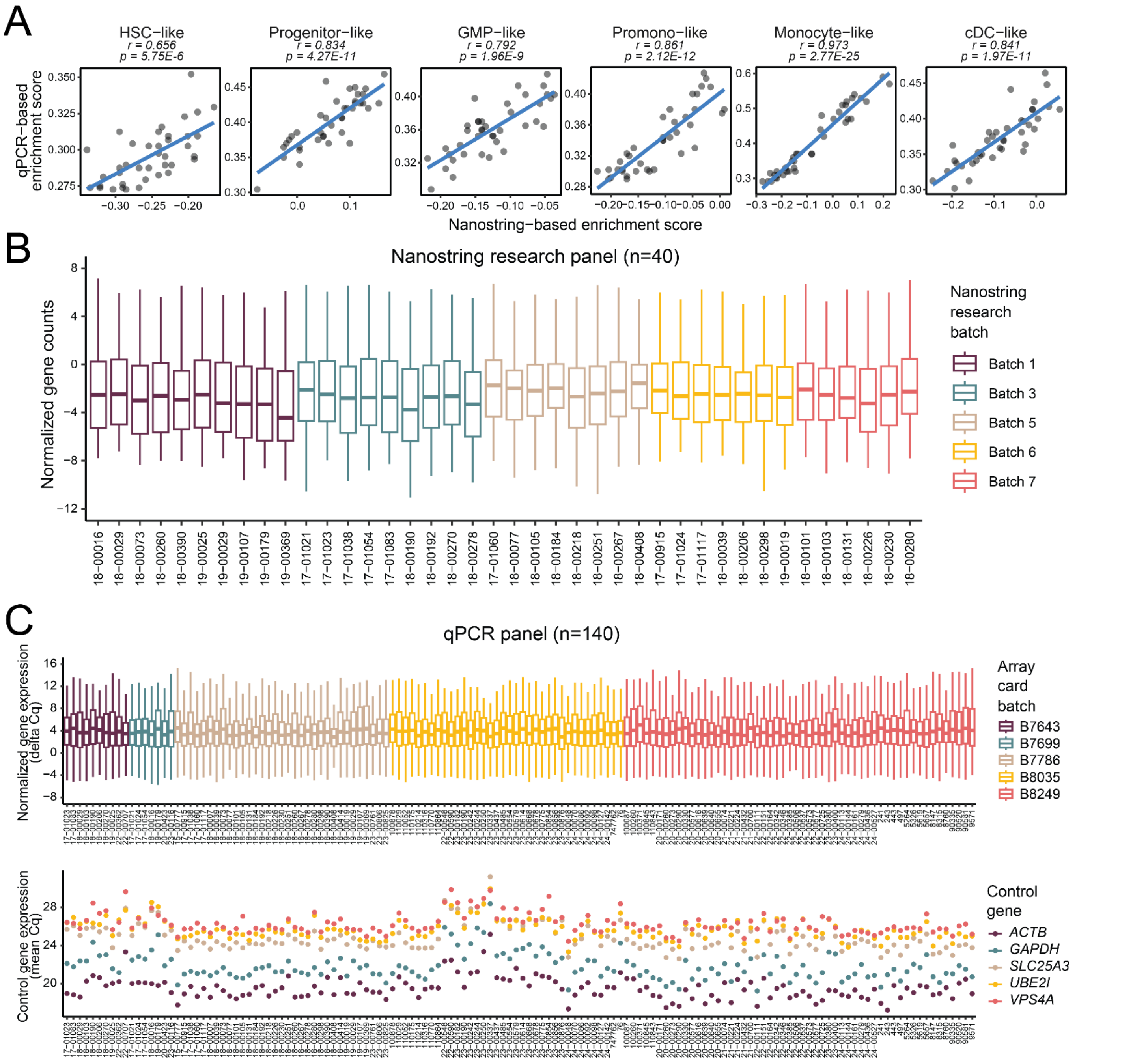


**Supplemental Figure S3. Consistency of research Nanostring- and qPCR-based gene expression distribution by sample and batch. (A)** Concordance of cell state stingscores with qPCR versus Nanostring platforms. Pearson r values are indicated. (**B**) Distribution of normalized gene counts for 40 AML patient samples profiled with the research Nanostring panel. Samples are grouped according to cartridge batch, with boxplots highlighting the interquartile range of all genes for a given sample. **(C)** Distribution of normalized gene expression for 140 AML patient samples profiled using the qPCR TaqMan array card assay (upper panel). Samples are grouped according to reagent batch order of TaqMan cards. Expression (mean Cq) values for each of five control genes on the qPCR panel are shown for each of the same samples as above, in the same order from left to right (lower panel).

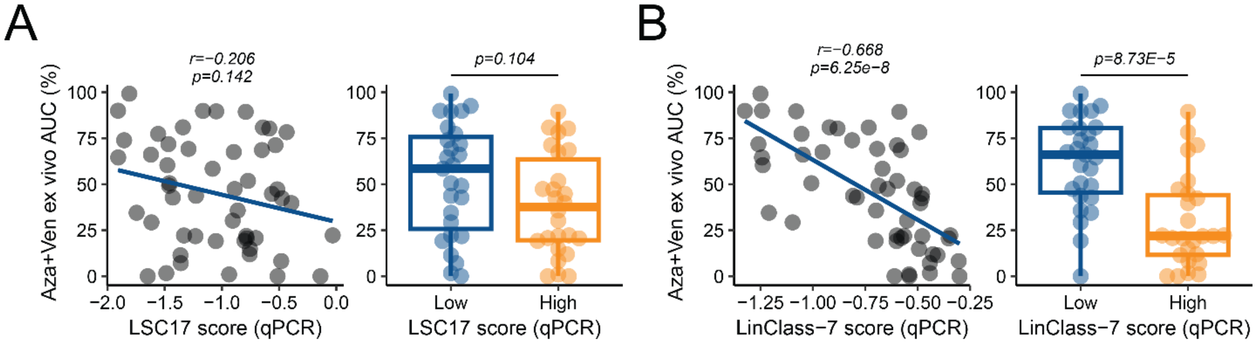


**Supplemental Figure S4. Correlation of qPCR-based stemness scores with ex vivo Aza+Ven sensitivity. (A)** Comparison of qPCR-derived scores for LSC17 versus ex vivo sensitivity to HMA+Ven for ND-AML samples (n=100). The median cut point for High vs Low LSC17 scores was established in Fig. 1B. **(B)** Comparison of qPCR-derived scores for LinClass-7 versus sensitivity to ex vivo HMA+Ven for ND-AML samples (n=100). The median cut point for High vs Low LinClass-7 scores was determined using the median score, as in [20].
